# A novel surfactant protein is associated with extrapulmonary respiration in lungless salamanders

**DOI:** 10.1101/261412

**Authors:** Zachary R. Lewis, Jorge A. Dorantes, James Hanken

**Author notes:** Present address: Department of Ecology and Evolutionary Biology, Yale University, New Haven, CT, USA, 06511. Present address: Universidad Autónoma de Guadalajara, Facultad de Medicina, Guadalajara, Jalisco, C.P. 44100, México. To whom correspondence should be addressed, Address: 165 Prospect St, New Haven, CT, 06511.

## Abstract

Numerous physiological and morphological adaptations were achieved during the transition to lungless respiration following evolutionary lung loss in plethodontid salamanders, including those that enable efficient gas exchange across extrapulmonary tissue. However, the molecular basis of these adaptations is unknown. Here we show that lungless salamanders express in the skin and buccal cavity—the principal sites of respiratory gas exchange in these species—a novel paralog of the gene Surfactant-Associated Protein C (SFTPC), which is a critical component of pulmonary surfactant expressed exclusively in the lung in other vertebrates. The paralogous gene appears to be found only in salamanders, but, similar to SFTPC, in lunged salamanders it is expressed only in the lung. This heterotopic gene expression, combined with predictions from structural modeling and respiratory tissue ultrastructure, suggest that lungless salamanders produce pulmonary surfactant-like secretions outside the lungs and that the novel paralog of SFTPC might facilitate extrapulmonary respiration in the absence of lungs. Heterotopic expression of the SFTPC paralog may have contributed to the remarkable evolutionary radiation of lungless salamanders, which account for more than two thirds of urodele species alive today.

## Introduction

Most amphibians must confront the challenges of respiring both in water and on land. To do so, they utilize numerous gas exchange surfaces including the lungs, gills, integument and buccopharyngeal mucosa, which are employed to varying extents depending on species and life-history stage. In adult salamanders, for example, the integument may be responsible for 50% or more of oxygen uptake [1,2]. Lability in sites of gas exchange is especially critical for metamorphosing species, which face different respiratory demands in air and water. Little is known, however, about the molecular mechanisms that enable the ontogenetic and evolutionary transitions from aquatic to aerial respiration. The mechanism of aerial respiration is even more enigmatic in lungless species, which rely entirely on extrapulmonary sites of respiration [1–3]. The family Plethodontidae includes more than two thirds of all living salamander species; most are fully terrestrial, and all adults lack lungs. Respiration takes place solely across the integument and buccopharyngeal mucosa, and also across the gills in aquatic larval forms, when present. Lunglessness is not unique to plethodontids—it has evolved several times in other amphibians, including salamanders, frogs and caecilians [4]—but its adaptive significance is unresolved [5,6].

How lungless salamanders are able to satisfy metabolic demands for oxygen is a topic of considerable interest. In theory, lunglessness limits thermal tolerance and maximum body size, yet lungless salamanders paradoxically occupy diverse thermal environments and attain relatively large body sizes. Plethodontids possess highly vascularized skin and buccopharyngeal mucosa, which may compensate for the loss of pulmonary respiration [1,7–9]. The buccopharyngeal membranes, in particular, may function as an adaptive respiratory surface that facilitates gas exchange, as evidenced by increased oscillation of the floor of the buccal cavity under hypoxia, high temperature, or activity, which presumably serves to draw more air into the mouth [1,7,10]. Selection for efficient extrapulmonary respiration may have played a major role in the adaptive radiation of terrestrial plethodontids [1]. Indeed, the evolution of highly efficient cutaneous and buccopharyngeal respiration is believed to have freed plethodontids from the ontogenetic and functional constraints associated with the use of a buccal pump for pulmonary ventilation, thereby enabling them to occupy diverse habitats and evolve ballistic tongue projection [2,11,12].

To identify the molecular adaptations that might facilitate lungless respiration, we investigated the expression of a crucial pulmonary surfactant-associated protein in plethodontid salamanders. Proper lung function requires pulmonary surfactant, a complex and evolutionarily variable mix of molecules that facilitate mucous spreading and lung compliance and improve oxygen diffusion [13–15]. Surfactant-associated protein C (SFTPC) is a hydrophobic protein found in pulmonary surfactant that localizes to the lung’s air-liquid interface. It reduces surface tension by aiding the adsorption and distribution of lipids within pulmonary surfactant and specifically enhances oxygen diffusion [14,16,17]. SFTPC also regulates production and turnover of phosphatidylcholine, a major component of pulmonary surfactant, and it limits the thickness of the hypophase, the liquid layer that lines the lung’s inner surface [18–20]. Mucous layer thickness greatly impacts gas exchange between the environment and the blood supply [21]. Additionally, oxygen uptake is enhanced by the presence of surfactant in the hypophase, likely due to convective effects that facilitate mixing of oxygen and mucous or increased rate of oxygen trafficking [15,22]. The expression of SFTPC is highly conserved among tetrapods: all species evaluated previously express SFTPC exclusively in the lungs [23–25] (and Supplementary Text).

## Results and Discussion

Despite lacking lungs as adults, plethodontid salamanders express SFTPC (Fig. S1). Due to the consistent restriction of SFTPC expression to the lungs and to SFTPC’s conserved role across tetrapods, we compared SFTPC expression between lungless and lunged salamanders. Surprisingly, several species of salamanders express two transcripts with high sequence identity to annotated SFTPC sequences (Fig. S1a, b). Both transcripts exclusively match SFTPC within the NCBI nucleotide collection database, but one possesses higher sequence similarity to amniote and frog SFTPC. We denote the transcript with lower similarity SFTPC-like. We did not find SFTPC-like within vertebrate genomes or transcriptomes outside of salamanders. Phylogenetic analyses support the hypothesis that SFTPC-like represents a previously undescribed salamander-specific paralog of the highly conserved lung-specific gene, SFTPC (Figs. S1, S3; Supplementary Text). SFTPC-like may maintain or partially maintain the characteristic hydrophobic alpha-helical configuration of the SFTPC mature peptide [16] (Fig. S1c).

Numerous studies of tetrapods localize SFTPC exclusively to the lungs [23–25]. SFTPC and SFTPC-like in the lunged salamander *Ambystoma mexicanum* match this highly conserved pattern (Fig. 1). As visualized by *in situ* hybridization, SFTPC and SFTPC-like staining is only observed in the extremely squamous alveolar epithelial cells lining the lungs and trachea (Figs. 1, S5, S6). SFTPC-like expression, however, is lower than SFTPC and the genes are expressed at different times: SFTPC-like is low or non-existent before hatching, whereas SFTPC is expressed beginning in embryos immediately following the formation of the laryngotracheal groove, a ventral outpocketing of the foregut that precedes lung outgrowth (Fig. 1a, b), and continuing into adulthood (Fig. 1d).

**Figure 1.**
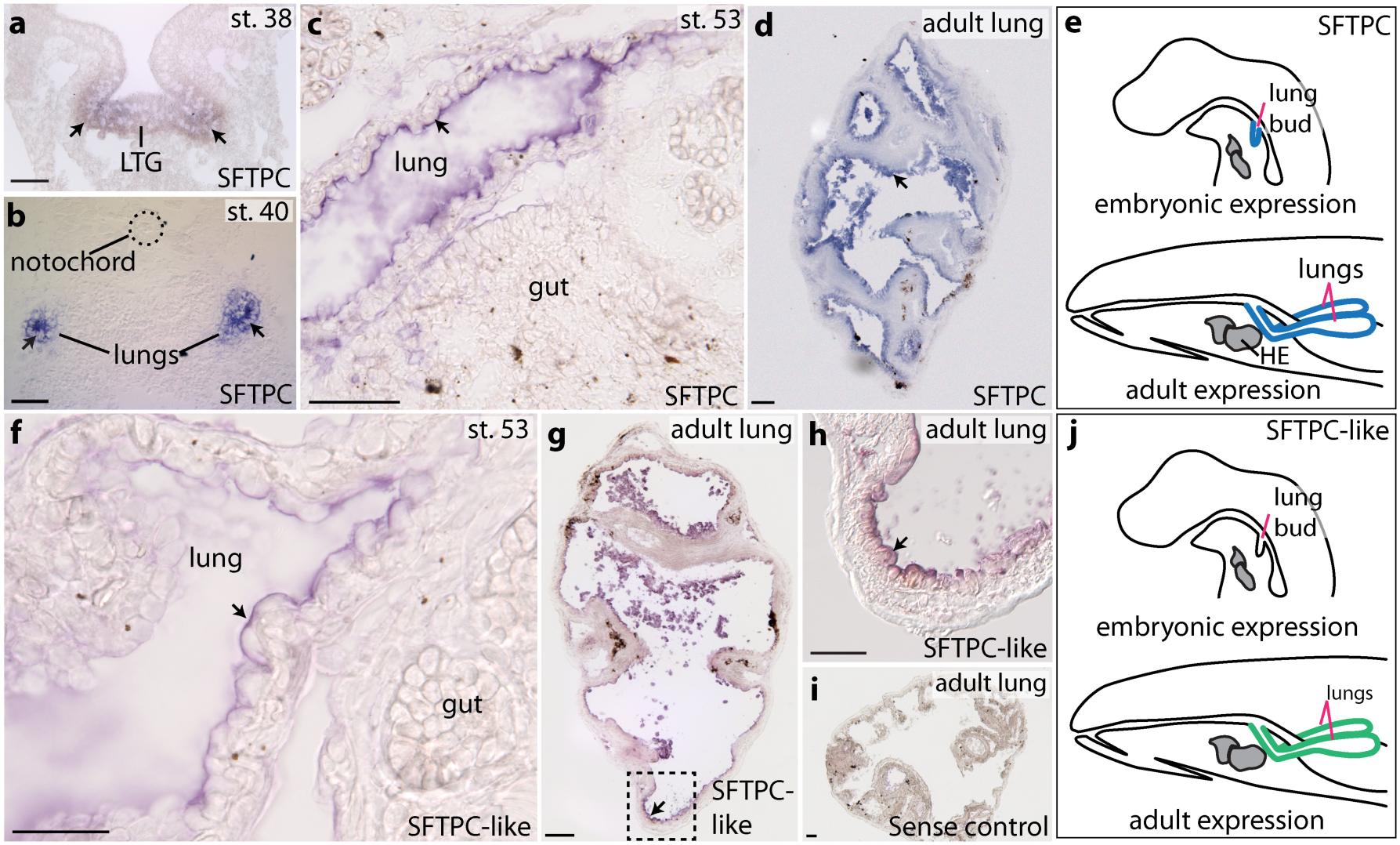
Expression patterns of Surfactant-Associated Protein C (SFTPC) and SFTPC-like in a lunged salamander, *Ambystoma mexicanum,* visualized with antisense wholemount *in-situ* hybridization. Arrows point to representative regions of expression. SFTPC is expressed in the embryonic laryngotracheal groove (LTG) **a,** or lungs **b–d,** of all stages examined between embryo (st. 40), juvenile (st. 53) and adult. **f–h,** SFTPC-like is also expressed specifically in the lung in juveniles and adults, but at a lower level than SFTPC; it is not expressed in embryos. **h,** Boxed region in **g. i,** Negative (sense) control run in parallel with SFTPC-like. **e, j,** Schematic sagittal views summarize the expression sites of SFTPC and SFTPC-like. Blue regions denote high expression; green indicates lower level. **a, b, d** and **g–i** depict transverse sections; **c** and **f** are sagittal sections, anterior to the left. Scale bars: 100 μm. Additional abbreviation: HE, heart.

In contrast, SFTPC-like is expressed dynamically in lungless salamanders. In embryos and early larvae of *Desmognathus fuscus*, a metamorphosing species, SFTPC-like is expressed throughout much of the integument, with reduced staining on the dorsal (internal) surface of the operculum (gill covering) and in the limbs (Fig. 2a-h). Expression begins to diminish in the integument immediately before metamorphosis, but at the same time it expands to the buccopharyngeal mucosa (oral epithelium) (Fig. 2i–l). Integumentary expression at this stage is patchy: remaining SFTPC-like-positive cells are displaced towards the apical surface and display an irregular morphology (Fig. 2i, k). Cessation of integumentary expression of SFTPC-like coincides with several metamorphic transitions, but especially molting [26], when the integument is profoundly remodeled from a simple stratified epithelium to a thickened pseudostratified tissue rich in acinous glands (Fig. S4, Fig. 2i, k).

**Figure 2.**
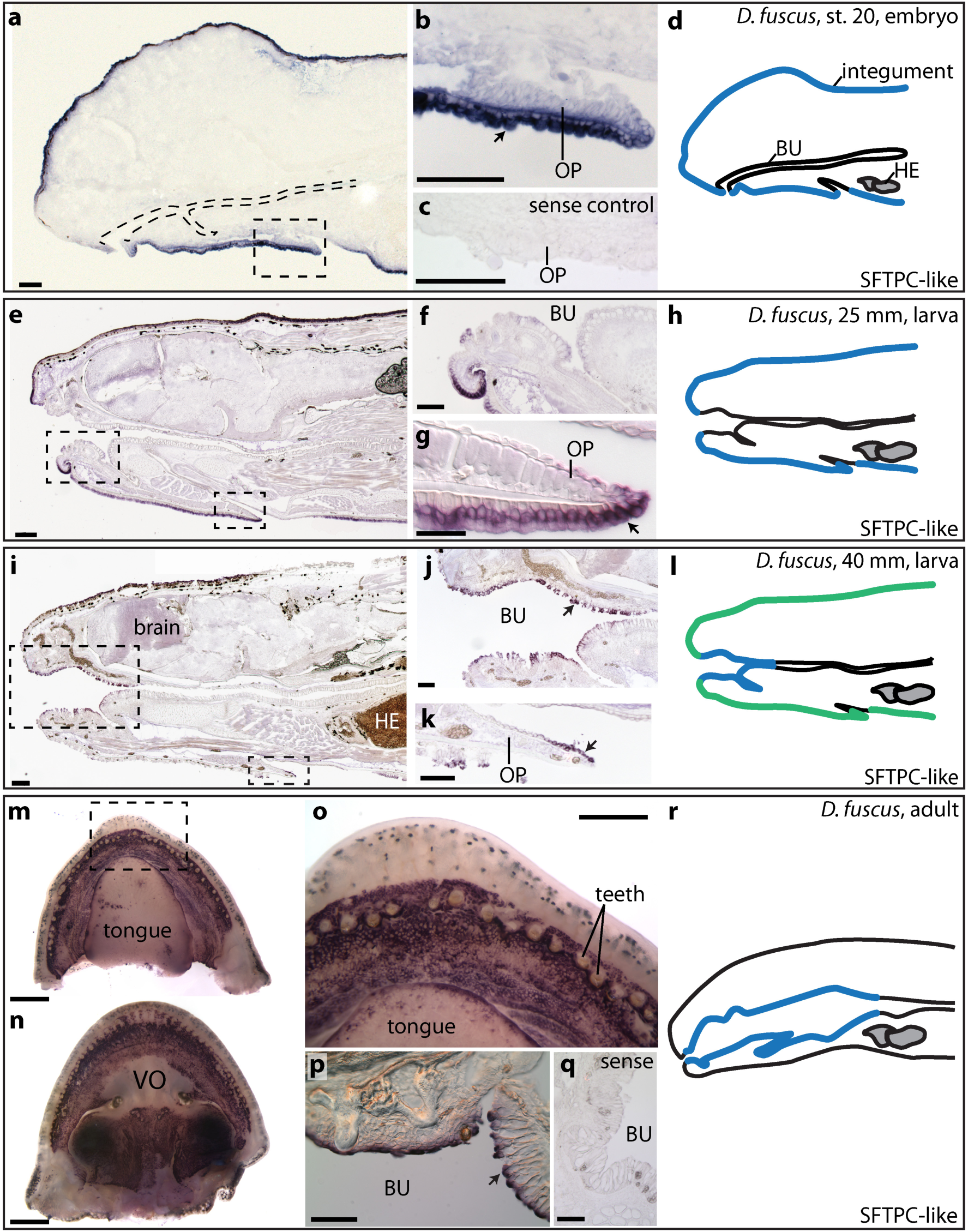
Expression of Surfactant-Associated Protein C-like (SFTPC-like) in the lungless salamander *Desmognathus fuscus,* visualized with antisense wholemount *in-situ* hybridization. Arrows point to representative regions of expression. **a–c,** In embryos, SFTPC-like is expressed in the integument. **b,** Boxed region in **a. c,** Negative (sense) control. **e–g,** SFTPC-like is expressed in the larval integument, 25 mm total length. **i–k,** SFTPC-like in a larva just prior to metamorphosis, 40 mm total length. Expression has declined in the integument but is now present in the buccopharyngeal mucosa (BU). **m–q,** Adult expression of SFTPC-like is confined to the buccopharyngeal cavity. **d, h, l, r,** Schematic sagittal views summarize the expression sites of SFTPC-like. Blue regions correspond to high expression; green indicates lower level. **a-l** are sagittal sections, anterior to the left; **p** and **q** are transverse sections. Scale bars: **a–c, e, f, i–k, p, q,** 100 μm; **g,** 50 μm; **m, n,** 1 mm; **o,** 0.5 mm. Additional abbreviations: HE, heart; OP, opercular covering; VO, vomer.

Immediately following metamorphosis, expression is absent from the integument and restricted to the buccopharyngeal mucosa (Fig. 2m–r). In adults, SFTPC-like is expressed in the buccal cavity and adjacent pharynx (Fig. 2m–r): it is confined to oral mucosa, with a strict boundary at the marginal teeth. SFTPC-like is not expressed along the vomerine bones (near the internal nares) or in the dorsal midline of the tongue, although it is strongly expressed along the tongue margin (Fig. 2m, o). Additional data from immunohistochemistry and mass spectroscopy are needed to determine if SFTPC-like transcripts are translated and whether this protein is then processed and secreted in a similar fashion to SFTPC in the lung.

Unlike SFTPC-like, SFTPC is expressed at an extremely low level in embryos of lungless species, which develop a transient lung rudiment [27]. The SFTPC transcript detected from transcriptome sequencing of the lung rudiment of *Plethodon cinereus* (Supplemental Data File 1) could not be cloned from either *P. cinereus* or *D. fuscus,* nor was it found in adult plethodontid transcriptomes. This suggests that SFTPC expression is inhibited by lung loss in lungless species, with the exception of SFTPC expression at certain embryonic stages in the presumptive lung region.

The larval integument of *Desmognathus fuscus* displays pronounced secretory activity (Fig. 3a–d). The outer layer—the stratum corneum—is a protective layer composed of anuclear keratinized cells. It displays an extremely high level of secretory activity evidenced by the near universal distribution of bilamellar secretory vesicles along its superficial surface (Fig. 3a, “SV”). Just basal to the stratum corneum is a layer of secretory cells, which are heavily vacuolated at their apical extent. The stratum corneum and the layer of secretory cells are not separated by a cell membrane, but a dark and consistent division appears between them (Fig. 3b). Bilamellar secretory vesicles virtually identical to those observed on the stratum corneum of *D. fuscus* are secreted from alveolar epithelial cells in the lung of *Ambystoma mexicanum* but are not present in its integument (Fig. 3e, g, h; Fig. S6, “SV”).

**Figure 3.**
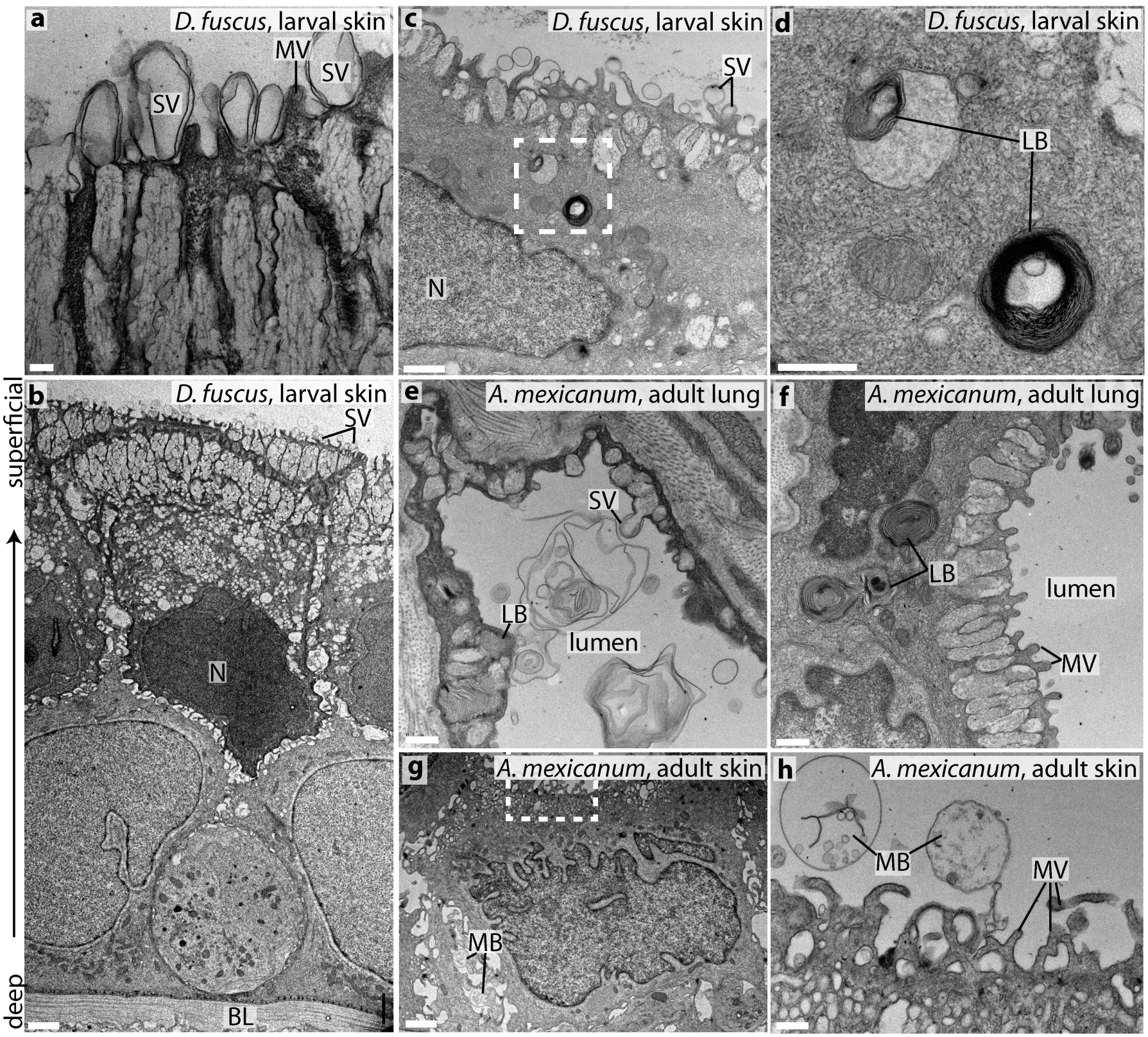
Secretory activity and lamellar bodies in the larval integument of lungless *Desmognathus fuscus* resemble those in the lung of *Ambystoma mexicanum*. **a–d**, Transmission electron micrographs of a 24-mm *D. fuscus.* **a,** The superficial (apical) surface is covered with secretory vesicles (SV), which emerge from columnar vacuolar structures, and is interspersed with microvilli (MV). **b,** Sagittal section through the epidermis; the superficial surface points upwards. **c,d,** Lamellar bodies (LB), indicative of surfactant production, are visible in the integument. The boxed region in **c** is enlarged in **d. e,f,** Lamellar bodies and secretory vesicles in the distal portion of the lung of an adult *A. mexicanum;* transverse section. **g,h,** Transverse section of gular integument of adult *A. mexicanum.* The boxed region in **g** is enlarged in **h.** Multivesicular bodies (MB) are visible in extracellular spaces in **g** and external to the skin in **h,** but there is no indication of active secretion of vesicles. The integument does not play a pronounced secretory role in this species. Additional abbreviations: BL, basal lamina; N, nucleus. Scale bars: **a,** 200 nm; **b, g,** 2 μm; **c,** 1 μm; **d, e, f, h,** 500 nm.

Pulmonary surfactant is trafficked in lamellar bodies in alveolar epithelial cells. These lamellar bodies are distinct from other ultrastructural lamellar elements in that they are large in diameter and spherical in shape [28–30]. Furthermore, these lamellar bodies are distinct from the bilamellar structures identified on the surface of the integument, in that they have many lamellae. The larval integument of *D. fuscus* contains large (0.5–0.75 μm diameter), spherical lamellar bodies (Fig. 3c, d, “LB”) that closely resemble those found in alveolar epithelial cells, which otherwise are known only from the lungs of other tetrapods.. Secretory activity and the presence of these distinctive lamellar bodies strongly suggest that *D. fuscus* produces surfactant in extrapulmonary sites of gas exchange, which correspond to sites of SFTPC-like expression.

Despite the lung- and trachea-specific expression of SFTPC and SFTPC-like in *A. mexicanum* revealed by *in situ* hybridization and the numerous reports of lung-specific expression of SFTPC in mammals (Supplementary text), recently published transcriptomes of *A. mexicanum* purport to map low numbers of SFTPC and SFTPC-like reads to several tissues, including blood vessels, bone, heart, regenerating limbs, and mixed stages of whole embryos [31]. Therefore, SFTPC and SFTPC-like may not be entirely lung-specific transcripts in lunged salamanders. Additional studies are needed, however, to evaluate the alternative interpretation that contamination, mapping or assembly issues might have yielded spuriously mapped reads.

Passive diffusion across a tissue layer, as occurs during cutaneous respiration, is a function of several variables. These include the size (area) of the surface over which gas exchange occurs and the thickness of the barrier between the underlying blood supply and the environment [3]. Barrier thickness depends mainly on the distance between the environment and the blood supply, but also on the thickness of the mucous layer between the environment and respiratory tissue. Diffusivity of mucus is about 30% lower than water [21]. Reduction of surface tension by pulmonary surfactant helps maintain a thin layer of airway surface liquid [20] and increases convection within the mucous, which together result in increased oxygen uptake [22]. SFTPC indirectly influences mucous layer thickness in the lung by regulating the production of phosphatidylcholine [18]. Pulmonary surfactant aids oxygen transport across the air-liquid interface, and hydrophobic surfactant proteins increase the rate of oxygen diffusion twofold over surfactant lipid alone [15,17]. SFTPC-like may facilitate respiration through a reduction of effective barrier thickness or an increase in diffusivity of the mucus layer. Pulmonary surfactant also plays non-respiratory roles, including facilitation of mucus spreading, innate immune defense, anti-edema agent, hydrostatic gas exchange, and preventing adhesion of lung surfaces when the lungs deflate [13,32,33]. It is possible that extrapulmonary surfactant produced in *Desmognathus fuscus* is performing one or more of these functions instead of, or in addition to, facilitating gas exchange.

Spatial and temporal expression of SFTPC-like and the ultrastructure of the associated integument suggest that, following SFTPC-gene duplication, SFTPC-like became neofunctionalized for extrapulmonary respiration in lungless salamanders. Sequence and structural conservation of SFTPC-like and SFTPC suggests that the two proteins function similarly (Fig. S1). However, to confirm neofunctionalization requires a more detailed functional characterization of SFTPC-like. This may include determining with increased phylogenetic and technical precision how expression of SFTPC-like has evolved in salamanders, proteomic characterization of skin and buccopharyngeal secretions, and assessment of whether SFTPC-like displays surface activity or aids gas exchange.

In metamorphosing species such as *Desmognathus fuscus,* respiratory and fluid-retention demands shift dramatically upon the transition from aquatic to terrestrial habitats [1]. Cutaneous water loss becomes a critical liability, but reduced skin permeability hinders cutaneous gas exchange. To counter this limitation, terrestrial plethodontids show increased reliance on buccopharyngeal respiration [1]. Ontogenetic shift of SFTPC-like from the skin to the buccopharyngeal cavity during metamorphosis (Fig. 2) correlates with the transition from aquatic to aerial respiration; SFTPC-like is expressed at the preferential sites of gas exchange at each life-history stage [1]. However, it is also possible that instead of playing a direct role in facilitating gas exchange, extrapulmonary surfactant balances fluid retention and respiratory demands by aiding fluid uptake from the mucus. Such a role would be consistent with the proposed anti-edema properties of surfactant within the lungs [13].

Gene duplication is increasingly recognized as a driving force of evolutionary innovation [34]. While gene duplication does not always lead to functional divergence [35,36], regulatory changes in duplicated genes, such as altered expression sites, may enable the evolution of novel traits in individual lineages. Many studies of the evolutionary phenomenon of adaptive radiation have emphasized morphological traits whose appearance in particular lineages promote phylogenetic and ecological diversification [37]. We propose that such morphological traits, or key adaptations, work in concert with novel and functionally significant molecular features to enhance evolutionary success, and that such instances of concerted evolution are more widespread than is currently recognized.

In plethodontid salamanders, it is possible that the combination of morphological adaptations [7,9,38] and novel deployment of a critical surfactant protein enables efficient extrapulmonary respiration via the buccopharynx and integument. Conserved expression of SFTPC-like in lunged salamanders relative to SFTPC may be due to dosage-sharing between SFTPC and SFTPC-like, which constrains tolerable mutations in SFTPC-like gene regulation [35]. Indeed, lung loss may have resulted in relaxed stabilizing selection for SFTPC-like gene regulation, thereby enabling the evolution of novel expression patterns. Greater understanding of the evolution and function of SFTPC-like in additional salamander species will yield a more complete picture of the evolution and consequences of lung loss, while functional studies of SFTPC-like promise to reveal whether this novel pulmonary surfactant protein plays similar roles to SFTPC or has potential therapeutic applications. Given the convergent evolution of lung loss in several amphibian lineages, it will be interesting to investigate the molecular physiology of other lungless taxa. Neofunctionalization of SFTPC-like may represent an additional mechanism by which plethodontid salamanders have become one of the most speciose and geographically widespread clades of vertebrates on earth, despite the theoretical limitations on thermal tolerance and body size imposed by lunglessness.

## Materials and Methods

### Animal husbandry

All animal protocols were reviewed and approved by Harvard’s Institutional Animal Care and use Committee (protocol 99-09). *Desmognathus fuscus* (Northern dusky salamander) embryos were field-collected from the following two localities under Massachusetts Department of Fish and Wildlife collection permits 080.11SCRA (2012), 027.13SCRA (2013), 083.14SCRA (2014), and 022.15SCRA (2015) and appropriate local permits: Ashfield, Mass. (42.483111, - 72.761263) and Mass Audubon Wachusett Meadows Preserve (42.450922, - 71.913009). Adults were collected from Willard Brook State Forest (42.671606, - 71.776156).*Plethodon cinereus* (Eastern red-backed salamander) embryos were field collected from Willard Brook State Forest (42.671606, -71.776156).

*Desmognathus fuscus* embryos were maintained at 15°C in 0.1x Marc’s Modified Ringer solution (MMR; 0.01 M NaCl, 0.2 mM KCl, 0.1 mM MgSO4, 0.2 mM CaCl2, 0.5 mM HEPES pH 7.4). Following hatching, larvae were fed *Artemia* spp. and maintained at 17–20°C until they metamorphosed at approximately seven months post-hatching. Older larvae were hand-fed blood worms. Embryos and larvae were sampled at intermediate stages from embryogenesis until 3–5 days post-metamorphosis and fixed overnight in MEMFA (0.1 M MOPS pH 7.4, 2 mM EGTA, 1 mM MgSO4, 3.7% formaldehyde) at 4°C, then dehydrated and stored at −20°C in 100% methanol. Adults were fixed in a similar manner immediately upon collection.

*Ambystoma mexicanum* (Mexican axolotl) embryos were obtained from the *Ambystoma* Genetic Stock Center, University of Kentucky, and maintained in 20% Holtfreter solution at 17°C. Larvae were raised similarly to larval *Desmognathus fuscus.* Fixation was performed as described above.

*Ambystoma mexicanum* were staged according to [39,40]. *Desmognathus fuscus* embryos were staged using a staging table derived for *Plethodon cinereus,* as their developmental timing and morphology are grossly similar [41].

### PCR

Embryonic cDNA from *Desmognathus fuscus, Plethodon cinereus,* and *Ambystoma mexicanum* was used to clone the gene SFTPC-like. RNA was isolated from homogenates of whole animals at a variety of embryonic stages using TRIzol Reagent (Invitrogen/Thermo Fisher Scientific, Grand Island, N.Y.) and reverse-transcribed to cDNA using iScript reverse transcriptase (BioRad, Hercules, Calif.). The gene SFTPC was cloned from *A. mexicanum* and repeated attempts were made to clone SFTPC from plethodontids. Degenerate and non-degenerate PCR primers were used (Table S1).

Primers were designed based on alignment of SFTPC sequences from *Desmognathus fuscus, Xenopus laevis, X. tropicalis, Anolis carolinensis, Neovison vison, Bos taurus, Monodelphis domesticus,* and *Homo sapiens.* All sequences but one were obtained from GenBank; the *D. fuscus* sequence was kindly provided by Dr. David Weisrock.

### Transcriptome assembly

Transcriptomes for *Plethodon cinereus* and *Ambystoma mexicanum* were prepared from microdissected tissue from pharyngeal endoderm and mesoderm of embryos and from the lung of a juvenile *A. mexicanum.* Total RNA was utilized for library preparation by using the IntegenX PrepX RNA-Seq Library Kit (IntegenX, Pleasanton, Calif.) on an Apollo 324 robotic sample preparation system (WaferGen Biosystems, Fremont, Calif.), closely following kit instructions. Agencourt Ampure XP beads were used for magnetic purification steps (Beckman Coulter, Indianapolis, Ind.). Beads were kept at room temperature for 15 min before starting block setup. Beads were added last to the block, after a 30-sec vortex to fully resuspend them.

Following cDNA synthesis, the concentration of samples was assessed by using a Qubit 1.0 fluorometer (Invitrogen/Thermo Fisher Scientific, Grand Island, N.Y.), high-sensitivity dsDNA reagents (Molecular Probes/Thermo Fisher Scientific, Grand Island, N.Y.), and Qubit Assay Tubes (Invitrogen). Samples were diluted to 20 μg/ml and then sheared on a Covaris S220 Focused ultrasonicator (Covaris, Woburn, Mass.) using a 72-sec protocol and targeting 220/320 bp output. TapeStation HS D1K tape (Agilent) was used to examine sheared DNA for optimal size range. BIOO Scientific NEXTflex DNA barcodes (#514102, Austin, Tex.) were diluted to 5 μm and used in the IntegenX PrepX DNA Library ILM prep kit (#P003279, Pleasanton, Calif.). Library prep was performed on the Apollo 324 using the kit manufacturer’s precise instructions. 5-μl aliquots of ~2-μg/ml samples were then subjected to four PCR amplification cycles by using NEB Next Master Mix (#M0541S, NEB, Ipswich, Mass.) and NEXTflex Primer Mix (BIOO Scientific) and the following cycle conditions: denaturation at 98°C for 120 sec; 5 cycles of 98°C for 30 sec, 6°C for 30 sec, and 72°C for 60 sec; and final extension for 5 min at 72°C. The Agilent Apollo 324 was used for cleanup of PCR samples using the built-in PCR cleanup protocol and Agencourt Ampure XP beads. Libraries were analyzed with Qubit 1.0, TapeStation and qPCR to assess library concentration, size and quality. Samples were each diluted to 0.29 nM concentration and then pooled. Two lanes of 2 x 150 bp Illumina HiSeq 2500 Rapid Run RNA-sequencing (Illumina, San Diego, Calif.) yielded a total 232.4 reads that passed filter.

Sequenced reads were trimmed with Trimmomatic [42] and concatenated. Ribosomal rRNA reads were removed by using Bowtie [43] and a custom database of known rRNA sequences for each species. Transcriptomes were assembled *de novo* using Trinity [44,45]. BLAST databases were created from the *de novo* assemblies and used to identify SFTPC and SFTPC-like sequences.

### SFTPC phylogeny

SFTPC sequences were identified through BLASTX, TBLASTN and TBLASTX searches of sequenced transcriptomes from this study (*Plethodon cinereus* and *Ambystoma mexicanum*) as well as from the transcriptomes listed in Table S2. Sequence identifiers and corresponding sequence data are provided as Supplemental Data File 1.

SFTPC sequences from annotated genomes were also taken from NCBI and ENSEMBL (Supplemental Data File 1). Outgroup proteins were selected based on previous phylogenies of SFTPC [46,47]. Predicted amino acid sequences were generated from all nucleotide sequences. Multiple sequence alignment was performed using PRANK [48]; resulting alignments were visually inspected (Supplemental Data File 2). ProtTest was used to identify an appropriate amino acid substitution model [49]. The optimal amino acid substitution model was JTT+G [50], as judged by AIC. Subsequently, 95% maximum clade credibility gene trees were reconstructed in MrBayes (v3.2.6) [51] using Markov chain Monte Carlo analysis with one million generations sampled every 100 generations and a relative burn-in of 25%. Convergence of the posterior probabilities was assessed by examining output statistics, including the potential scale reduction factor, which equaled or exceeded 1.000.

A phylogeny was also constructed in RAxML (v8.2.9) [52] by using 1000 bootstrap replicates and the aforementioned amino acid substitution model. Tree topology was concordant with the Bayesian tree generated in Mr. Bayes. JalView was used to generate the multiple sequence alignment image [53].PHYLDOG [54] was used to estimate gene duplication of SFTPC. A guide tree was constructed using NCBI taxonomy for major groups and [55] for amphibian relationships (Fig. S2). Topology optimization was not used.

### *In situ* hybridization

Embryos were fixed overnight in 4% paraformaldehyde (PFA) or MEMFA at 4°C, dehydrated and stored in 70% or 100% MeOH at −20°C. Wholemount mRNA *in situ* hybridization (ISH) was performed by rehydrating samples, which were then treated with 5–10 μg/ml proteinase K for 30–60 min, washed with PBTw (137 mM NaCl, 2.7 mM KCl, 10 mM Na2HPO4, 1.8 mM KH_2_PO_4_ and 0.2% Tween-20), post-fixed in 4% PFA, washed with PBTw, and pre-hybridized in hybridization buffer for 2 hr at 65°C (hybridization buffer: 50% formamide, 5x SSC, 0.1 mg/ml heparin, 1x Denhardt solution, 0.01% CHAPS, 0.2 mg/ml tRNA and 0.1% Tween-20; all solutions were RNase-free). DIG-labeled riboprobes were diluted approximately 1:40 in hybridization buffer, denatured at 85°C for 10 min, and then added to specimens. Hybridization was carried out overnight at 65°C.Posthybridization washes were performed with a solution of 50% formamide, 5x SSC and 0.2% Tween-20 at 65°C for eight changes of 30 min each. Specimens were washed with maleic acid buffer plus 0.2% Tween-20 (MABT) prior to blocking and antibody incubation. Antibody block solution included 20% heat-inactivated sheep serum and 2% blocking reagent (Roche, Penzberg, Germany) in MABT. Samples were incubated overnight at 4°C with 1:2500 anti-DIG-AP Fab fragments (Roche) diluted in blocking solution. Extensive washes with MABT were performed prior to color development using BM-Purple (Roche) or NBT/BCIP (Sigma, St. Louis, Mo.). Color development occurred over several hours. Embryos were then embedded for cryosectioning at 14–16 μm thickness. Photographs were taken using a Leica DMRE microscope (Wetzlar, Germany) equipped with a QImaging Retiga 2000r camera and a QImaging RGB slider (Model: RGB-HM-S-IR; Surrey, Canada) and Volocity 6.0 software (PerkinElmer, Waltham, Mass.).

### Structure models

An experimentally determined protein data bank (PDB) model for porcine SFTPC [16] was downloaded from the Research Collaboratory for Structural Bioinformatics PDB. The secondary and supersecondary structure of *Desmognathus fuscus* SFTPC-like was predicted using Quark *Ab initio* Protein Structure Prediction [56]. The N- and C-terminal extent of the *Desmognathus* SFTPC-like sequence was chosen based on alignment with mature forms of SFTPC found in mammals. Protein structure prediction was also performed with I-TASSER [57].

A structure model for *D. fuscus* SFTPC-like was also predicted in SWISS-MODEL [58] using the molecular structure derived by Johansson et al. [16] as a template. The PDB files for each SFTPC-like model were imported into PyMol (Schrödinger) and aligned with the SFTPC model to graphically illustrate structural similarities.

### Transmission electron microscopy

Two 24-mm (total length) *Desmognathus fuscus* larvae were euthanized and decapitated. Specimens were then dissected in fixative (2.5% glutaraldehyde and 2% paraformaldehyde in 0.1 M HEPES; the aldehydes were free of alcohol stabilizers). The head was cut into three 1-mm sagittal sections. An 18-cm adult *Ambystoma mexicanum* was euthanized and then dissected in fixative. Samples of the gular skin from the ventral head, the oral epithelium and the lungs were trimmed to 1-mm-thick pieces in fixative and fixed as above.

The samples were left in fixative for 3 d and then washed twice quickly with 0.1 M HEPES and three times for 5 min each with Milli-Q H_2_O (mqH_2_O). Next, samples were fixed for 24 hr at 4°C in aqueous 1% osmium tetroxide, followed by five washes in mqH_2_O for 5 min each. Subsequently, specimens were stained with 2% uranyl acetate (EMS, Hatfield, Pa.) overnight at 4°C, then washed two times for 5 min each with mqH_2_O. Specimens were dehydrated with 5-min washes of 50%, 70% and 95% ethanol, followed by three 10-min washes with 100% ethanol, then two quick rinses with propylene oxide (PO). Specimens were embedded in Embed 812 resin (EMS) formulated to medium hardness by rinsing 30 min each in 1:1 PO to Embed 812, 1:2 PO to Embed 812, then 60 min in 1:4 PO to Embed 812. Specimens were then transferred to 100% Embed 812 and incubated overnight at room temperature, followed by two subsequent changes of Embed 812, over a total embedding time of 48 hr. Samples were then positioned in molds and placed at 60°C for 3 d to polymerize.

Sectioning was performed on a Leica UCT ultramicrotome, using glass knives for trimming blocks and generating semi-thin (1 μm) sections, and a DiATOME diamond knife for generating thin sections of approximately 60–100 nm thickness (target thickness: 80 nm). Sections were flattened with chloroform vapor, transferred onto precoated Formvar/carbon 200 mesh copper grids (#01803F, Ted Pella, Redding, Calif.), and dried on filter paper.

Grids were imaged with an FEI Tecnai G2 series F20 transmission electron microscope (Hillsboro, Ore.) at 80 kV using a Gatan CCD camera and Gatan Digital Micrograph Software (Pleasanton, Calif.).

## Data Availability

All data are available as Supplemental Data Files 1 and 2.

## Acknowledgements

Sequences were provided by Chris Amemiya, John Burns, Paul Hime, Ryan Kerney, Justin Kratovil, Rachel Mueller, Igor Schneider, Randal Voss, David Weisrock, and Ryan Woodcock (Table S2). Carolyn Marks offered help and guidance with TEM. Carolyn Eng assisted with field collection. This work was performed in part at the Harvard University Center for Nanoscale Systems (CNS), a member of the National Nanotechnology Coordinated Infrastructure Network (NNCI), supported under NSF ECCS award no. 1541959. Z.R.L. was supported by the NSF Graduate Research Fellowship Program.

## Author contributions

Z.R.L. cloned SFTPC and SFTPC-like from *A. mexicanum* and *P. cinereus,* generated and analyzed transcriptomes, performed phylogenetic and structural analyses, performed TEM, collected and raised animals, generated gene expression data and participated in writing the manuscript. J.A.D. cloned SFTPC-like from *D. fuscus* and assisted with characterizing the expression of SFTPC-like. J.H. participated in the data analyses and writing of the manuscript.

## Competing interests

The authors declare that they have no competing financial interests.

## Supporting Information

### Supplemental Text

#### SFTPC expression specificity

The expression pattern of SFTPC is highly conserved: all tetrapods express SFTPC exclusively in the lungs [1–9]. In anamniotes, SFTPC is expressed throughout the lung [5,6,10], whereas in mammals it is confined to alveolar type II cells. Four reports cite expression of SFTPC outside of the lungs in humans, but each report has methodological problems, including possible contamination, that may make such claims unreliable. Mo et al. (2006) report SFTPC in human fetal and adult skin [11]. This claim, however, relies on immunohistochemical data obtained with an antibody that may yield spurious labeling, and on RT-PCR data that was not followed up with sequencing. RT-PCR is subject to contamination and mispriming. Bräuer et al. (2009) report SFTPC expression in submandibular and parotid glands based on RT-PCR and western immunoblots [12]. The western blots reveal a protein of the expected size of the SFTPC pro-protein, and RT-PCR of focal cDNA is performed alongside a positive control, but there is no follow-up sequencing. Schicht et al. (2015) report SFTPC in saliva from human patients based on western blots and ELISA [13]. The antibody used to detect SFTPC is not identified, however, and the isolated band is at 16 kDa while SFTPC proprotein is 21 kDa and mature SFTPC 3.7 kDa [14]. Additionally, saliva may be subject to contamination by surfactant produced in the lungs. Finally, Schob et al. (2013) report SFTPC expression in central nervous system tissue and cerebrospinal fluid based on RT-PCR, but they fail to rule out the possibility of genomic DNA contamination [15]. Their western blots also fail to demonstrate a band at the expected size for SFTPC given the antibody they employed, and the authors express confusion about how mRNA for SFTPC could be present in cerebrospinal fluid. In sum, there are problems with all recent studies that cite extrapulmonary expression of SFTPC in humans. At the same time, numerous studies in mammals and frogs, including ISH and reporter knockins, have failed to demonstrate extrapulmonary expression of SFTPC. For instance, a mouse line with Cre recombinase knocked in to the SFTPC locus fails to label cells outside of the lung when crossed to a reporter line [16]. Nevertheless, it remains possible that humans and perhaps other animals endogenously express SFTPC outside of the lungs. Optimally,*in situ* hybridization and transcriptome sequencing should be used to validate the human results presented above.

#### Evidence for Duplication of SFTPC

Both SFTPC and SFTPC-like diverge within exonic regions, but not according to putative splice boundaries (Fig. S1a), which indicates that SFTPC-like is not an isoform of SFTPC. While SFTPC-like is divergent from SFTPC sequences, it is not an ortholog of a closely related BRICHOS domain-containing gene (Fig. S1b). We found SFTPC-like expressed in ten species of lunged and lungless salamanders (Fig. S1a, b; Supplemental Data File 1), most of which also express SFTPC. SFTPC-like is not found in genomes or transcriptomes of any other tetrapods, or even other vertebrates.

The most parsimonious explanation for the corresponding gene tree is that the tetrapod ortholog of SFTPC was duplicated in the salamander lineage following its divergence from frogs, followed by substantial sequence divergence between SFTPC and SFTPC-like (Fig. S1b). Low statistical support at the SFTPC-like node should be interpreted as a polytomy, and long-branch attraction likely has caused an artifactual affiliation between coelacanth and lungfish SFTPC orthologs and salamander SFTPC-like. We applied PHYLDOG [17] to explicitly test for gene duplications of SFTPC. Given a guide tree with known phylogenetic relationships (Fig. S2), PHYLDOG predicts that SFTPC-like arose by gene duplication (Fig. S3). SFTPC-like has been meiotically mapped to linkage group 6 in *Ambystoma mexicanum,* a lunged salamander, and is located within a region syntenic to human chromosome 15 [18]. SFTPC-like and SFTPC have been assembled to two separate genomic scaffolds from *A. mexicanum* [19], supporting SFTPC-like’s origin via gene duplication.

## Supporting Information

**Table S1.**
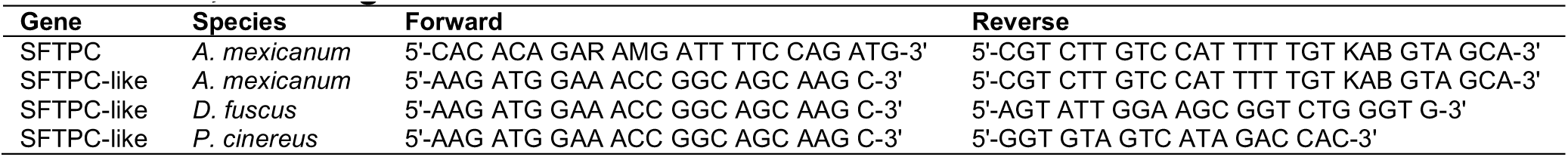
Primers used to clone SFTPC and SFTPC-like from *Ambystoma mexicanum, Desmognathus fuscus* and *Plethodon cinereus.*

**Table S2.** Transcriptomes used to identify SFTPC and SFTPC-like.

| Species | Transcriptome Source |
| --- | --- |
| <u>Salamanders</u> |  |
| *Ambystoma andersoni* | Ryan Woodcock and Randal Voss |
| *Ambystoma mexicanum* | Present study |
| *Ambystoma tigrinum* | Ryan Woodcock and Randal Voss |
| *Ambystoma tigrinum* | [20] |
| *Cryptobranchus alleganiensis bishopi* | David Weisrock and Paul Hime |
| *Cynops cyanurus* | David Weisrock and Paul Hime |
| *Desmognathus fuscus* | David Weisrock and Justin Kratovil |
| *Ensatina eschscholtzii* | Rachel Mueller [21] |
| *Hynobius chinensis* | [22], reassembled by Paul Hime |
| *Hynobius retardatus* | [23] |
| *Notophthalmus viridescens* | [24] |
| *Plethodon cinereus* | Present study |
| <u>Frogs</u> |  |
| *Pelophylax nigromaculatus* | [25] |
| *Rana (Lithobates) pipiens* | [26] |
| <u>Dipnoi</u> |  |
| *Lepidosiren paradoxa* | Igor Schneider [27] |
| *Protopterus annectans* | Chris Amemiya [27] |

## Supporting Information Figures

**Figure S1.**
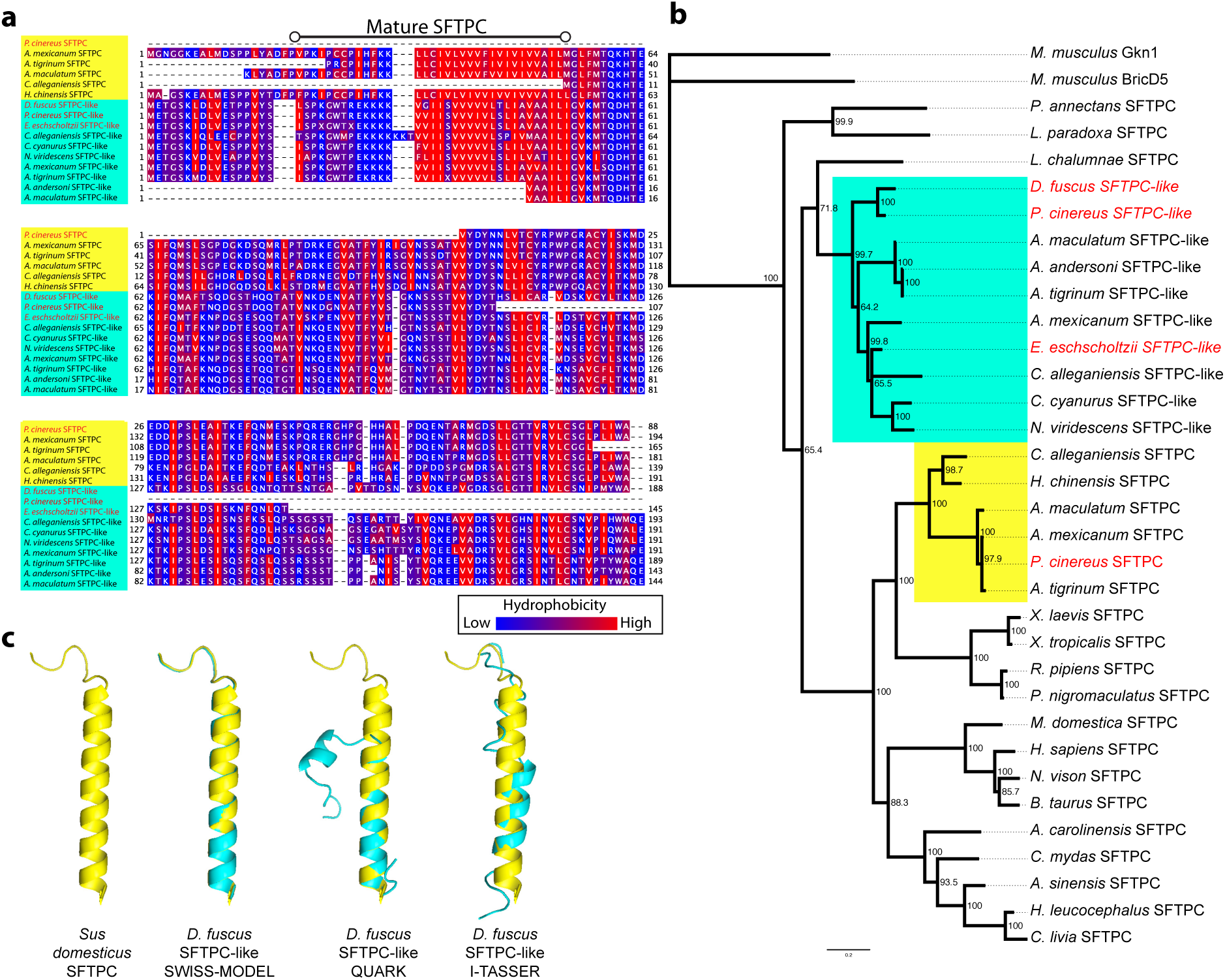
A novel form of Surfactant-Associated Protein C (SFTPC) is expressed in several species of salamanders. **a**, Amino acid alignment of SFTPC (yellow) and SFTPC-like (cyan) sequences reveals conservation of hydrophobic residues within the mature peptide domain. Full species names and accession numbers are listed in Supplemental Data File 1. Lungless (plethodontid) species are in red font. **b,** Bayesian 95% maximum clade credibility tree for SFTPC reveals SFTPC-like transcripts in 10 species of salamanders. SFTPC-like is not a related ortholog because it is nested within the SFTPC phylogeny. SFTPC-like is not found in any genome or transcriptome outside of salamanders. Node values are posterior probabilities; scale bar is expected changes per site. **c,** Predicted secondary structure of SFTPC-like from *Desmognathus fuscus.* SFTPC-like structure predictions (cyan) utilizing SWISS-MODEL [28], QUARK *Ab initio* predictions [29] and I-TASSER [30] are aligned with the resolved SFTPC mature peptide (yellow)[31].

**Figure S2.**
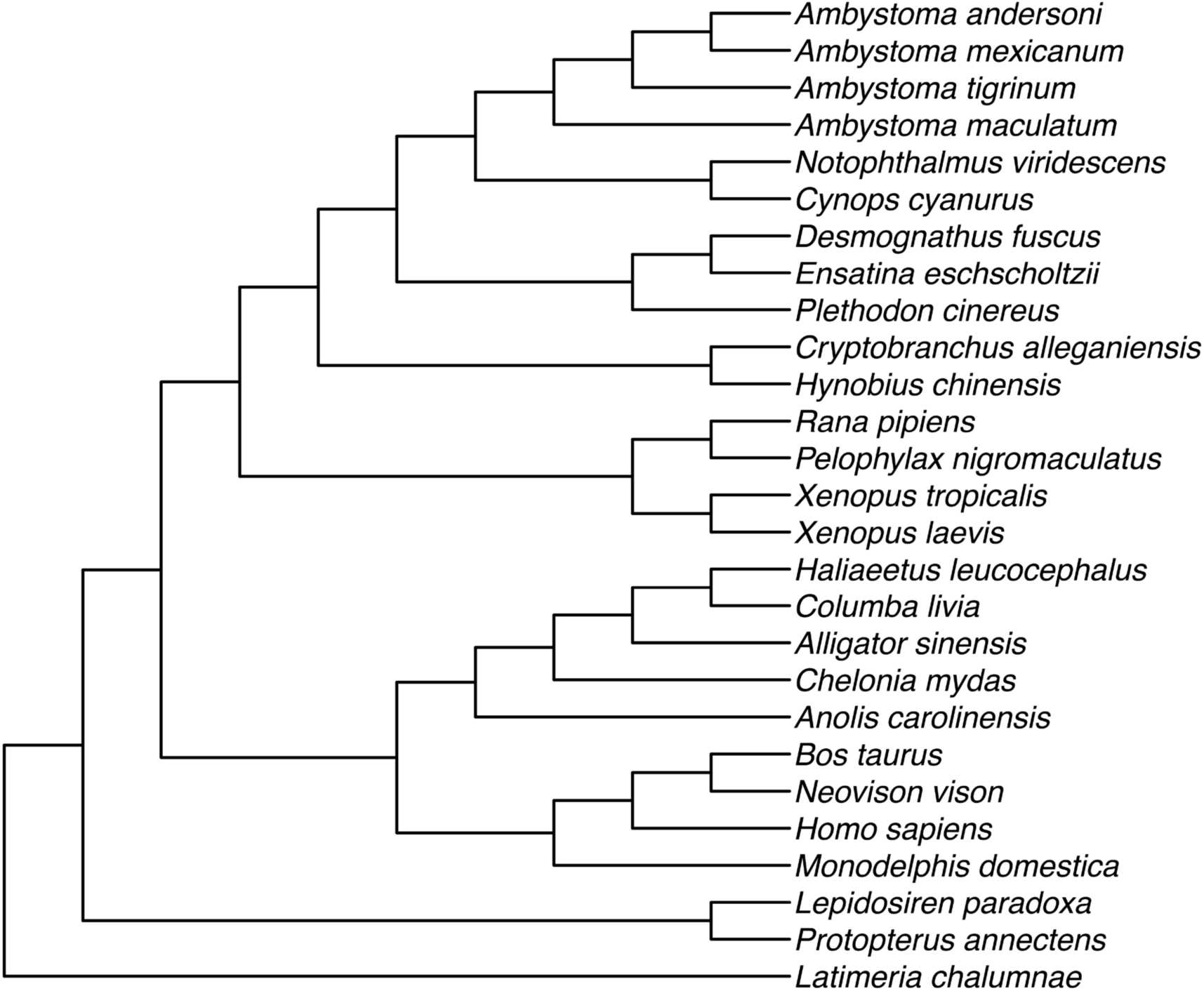
Guide tree for PHYLDOG. NCBI taxonomy was used to generate the tree, combined with Pyron and Wiens (2011) [32] for amphibian taxonomy.

**Figure S3.**
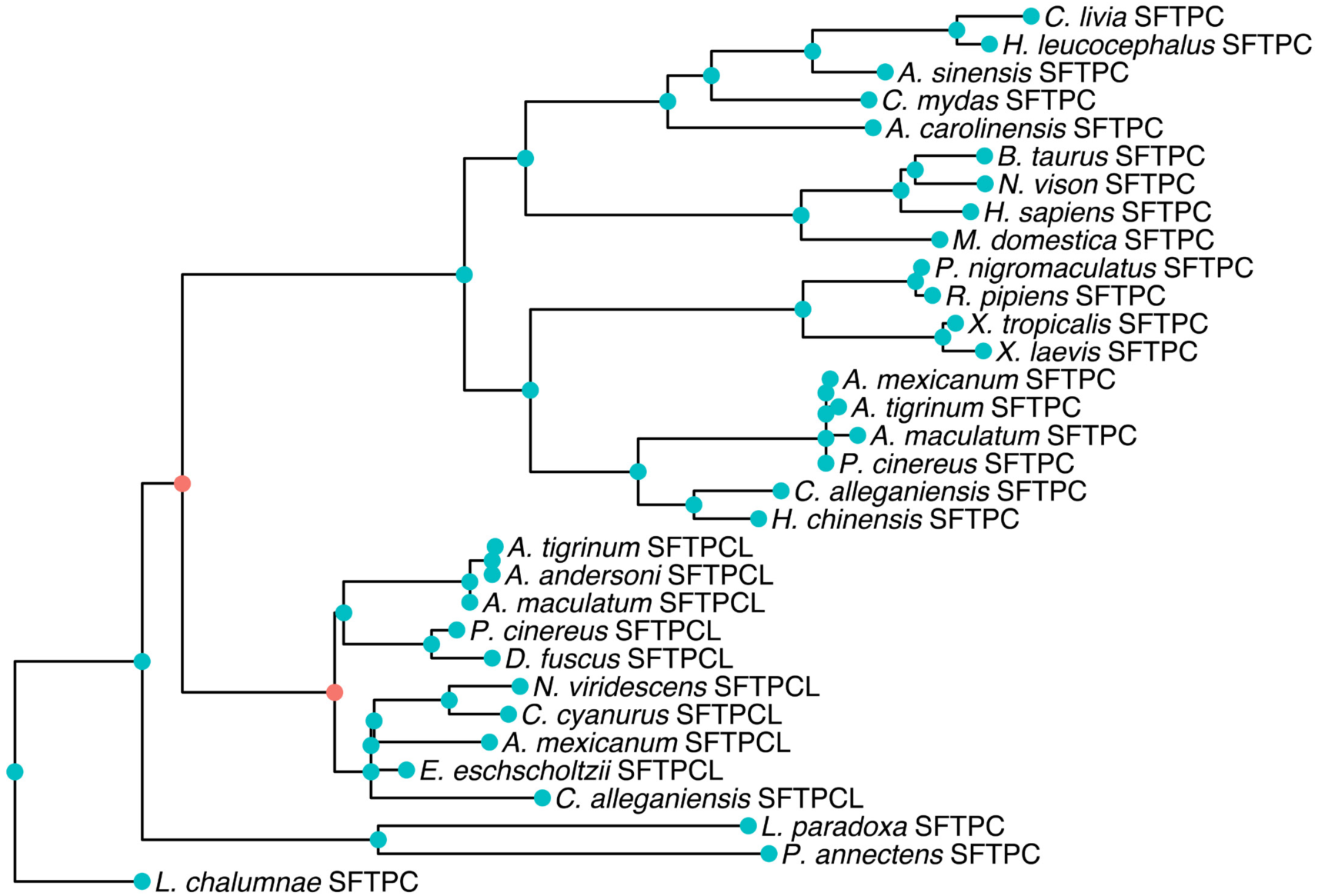
Gene duplications predicted by PHYLDOG. The two nodes where a gene duplication event is predicted are colored red. Blue nodes indicate divergence due to speciation events. PHYLDOG predicts that SFTPC-like (SFTPCL) originated due to gene duplication. A second duplication event of SFTPC-like is predicted in salamanders. While most species of salamanders appear to express only one form of SFTPC-like, several species express SFTPC-like transcripts with slight sequence differences, as noted in Supplemental Data File 1. Further work is needed to determine if these sequence differences can be attributed to further duplications of SFTPC-like or are the result of assembly error or alternative splicing. Only one sequence per species was selected for phylogenetic analysis. Complete species names are provided in Figure S2 and Table S2.

**Figure S4.**
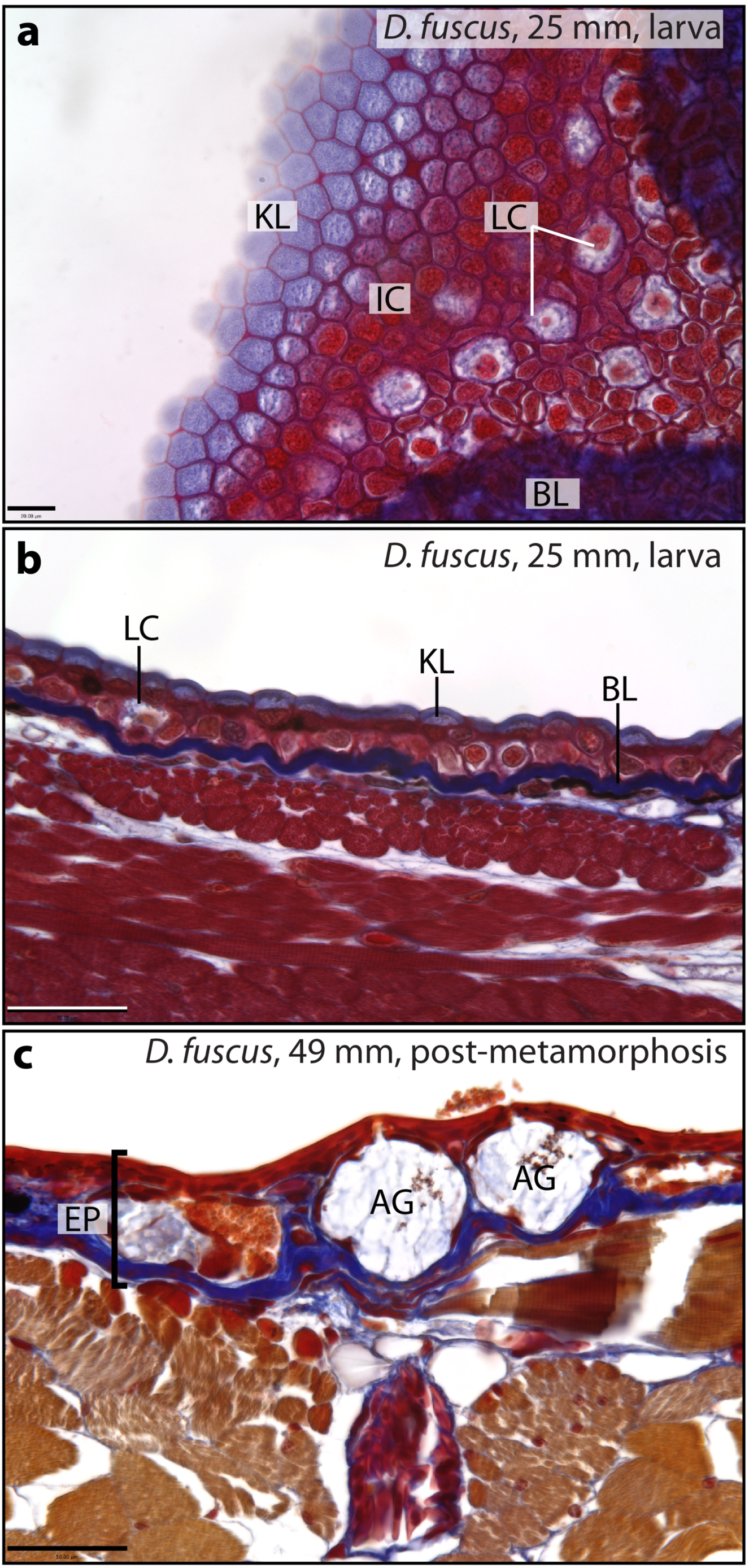
Histology of the integument in *Desmognathus fuscus* before and after metamorphosis. **a**, Tangential section through the gular region of a larva shows the layers of the integument (from left to right): flattened, cuticle-like keratinized layer (KL); an inner cell layer (IC); large cuboidal Leydig cells (LC) intermingled with capillaries and other supporting cells; basal lamina (BL). **b,** Sagittal section from the abdominal region of a larva. **c,** Transverse section from a recently metamorphosed specimen showing the acinous glands (AG) and a greatly thickened epidermis (EP). Mallory’s trichrome stain. Scale bars: **a,** 20 μm; **b,c,** 50 μm.

**Figure S5.**
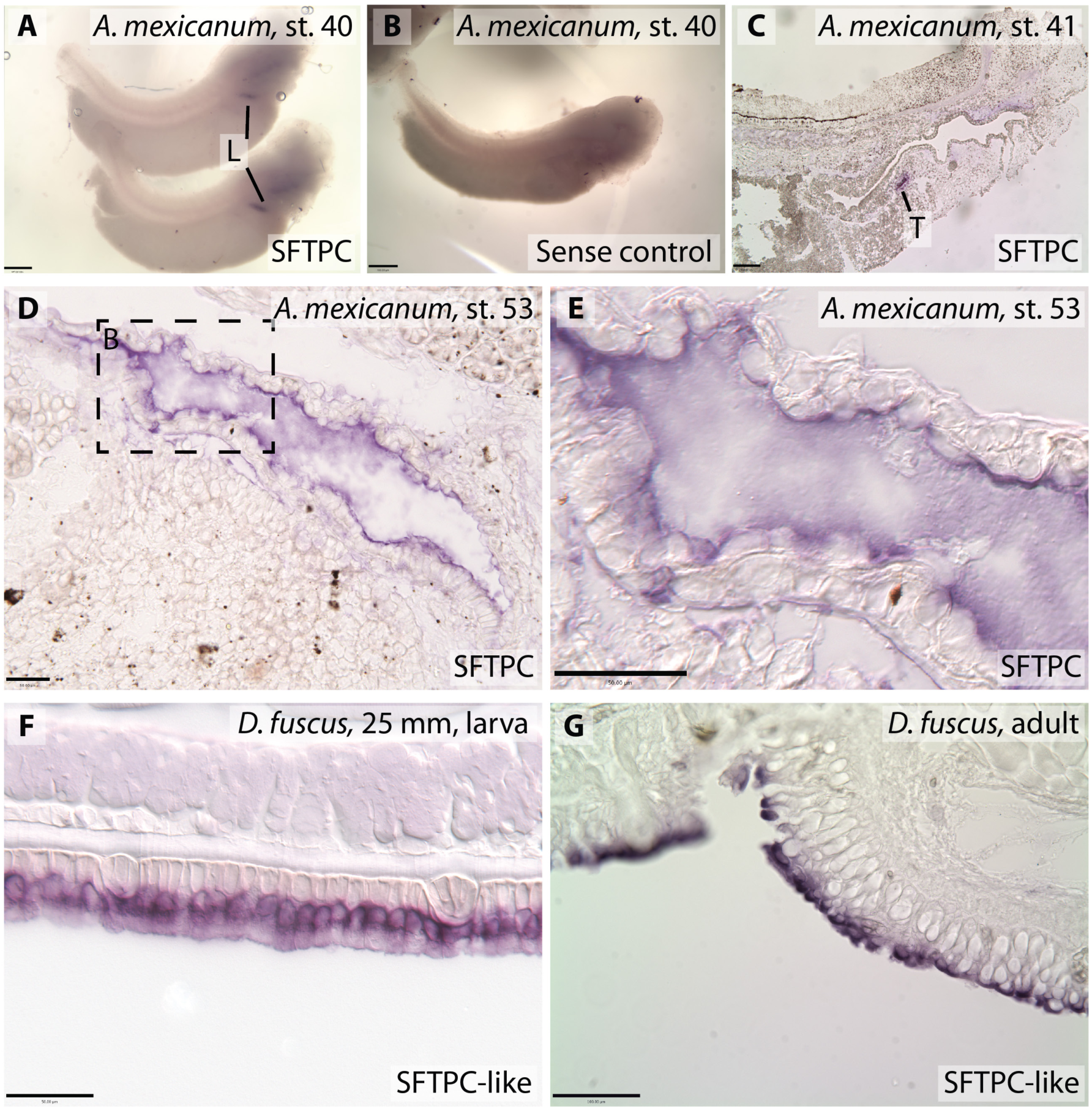
Additional images of SFTPC and SFTPC-like expression patterns. **a,** Wholemount embryos of *Ambystoma mexicanum* display SFTPC expression specific to the lungs (L). **b,** SFTPC sense control at the same stage shows no lung expression. **c,** Midsagittal section of *A. mexicanum* embryo stained for SFTPC shows expression in the trachea (T), but no expression in the integument or buccopharynx. **d,e,** SFTPC expression in *Ambystoma mexicanum* lung is confined to the squamous epithelial cells lining the lumen. **e** is an enlargement of the boxed region in **d. f,** SFTPC-like expression in *Desmognathus fuscus* integument is confined to the apical cellular layer. **g,** SFTPC-like expression in adult *D. fuscus* buccal epithelium. Scale bars: **d,e,f,** 50 μm; **a,b,c,g** 100 μm.

**Figure S6.**
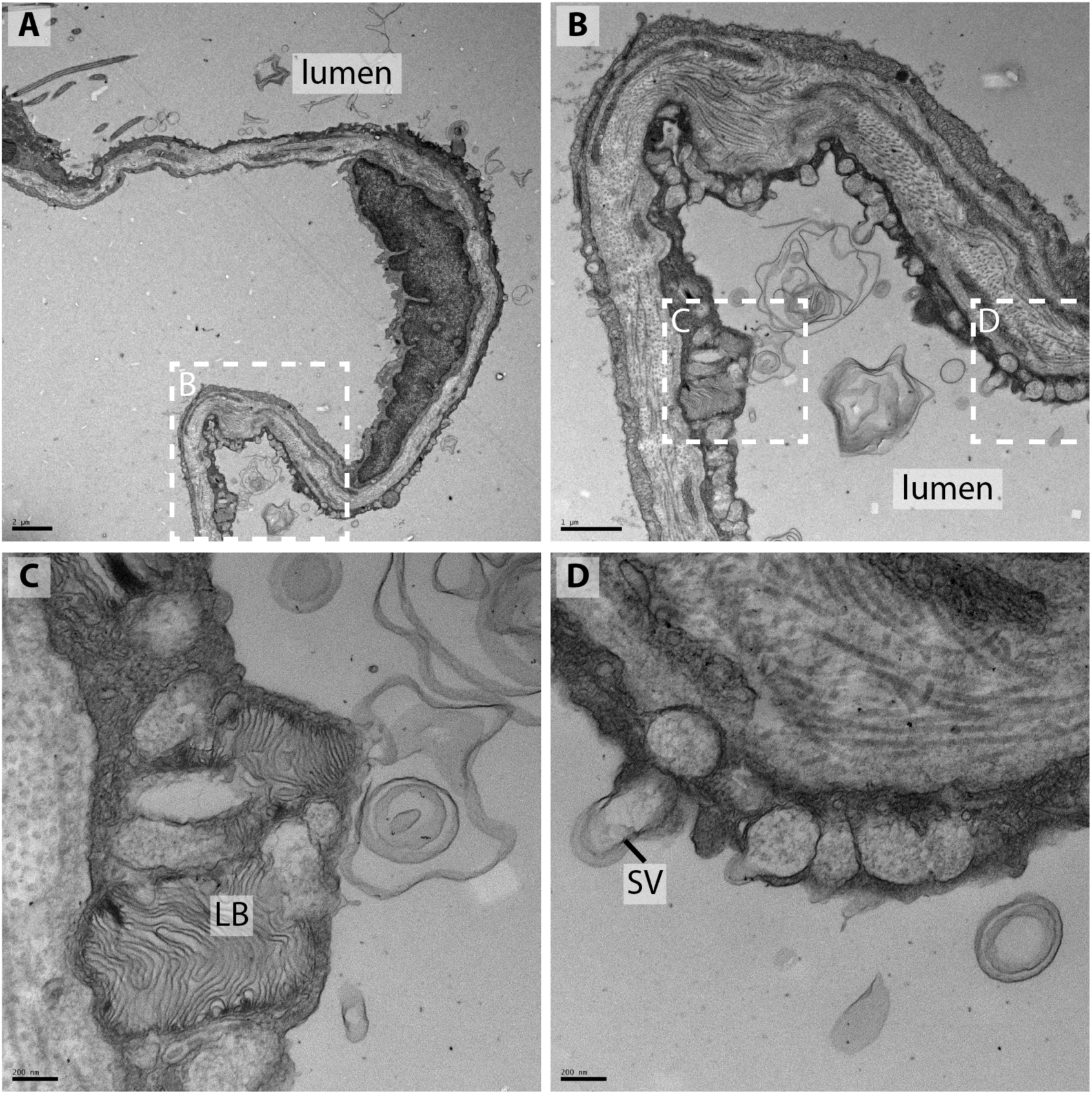
Ultrastructure of alveolar epithelial cells in adult *Ambystoma mexicanum.* **a,** low magnification view of the pulmonary epithelium. The lumen of the lung is to the right. **b,** enlargement of boxed area in **a. c,d,** enlargement of boxed regions in **b** show lamellar bodies (LB) and secretory vesicles (SV). Scale bars: **a,** 2 μm; **b,** 1 μm; **c,d,** 200 nm.

## Captions for Supplemental Data Files

Supplemental Data File 1: Excel spreadsheet with all sequence data used for the study.

Supplemental Data File 2: A FASTA amino acid alignment used to generate the gene tree.

